# Submembrane geography dictates GPCR signalling compartmentation and physiological function

**DOI:** 10.64898/2026.10.30.763044

**Authors:** Alexander Froese, Bing Qi, Miao Feng, Greta Böer, Karoline Morhenn, Svenja Kupsch, Xing Sun, Hariharan Subramanian, Jose L. Sanchez-Alonso, Roman Medvedev, Peter T. Wright, Marta Gallego, David Revuelta, Nadja I. Beck, Cristina E. Molina, Michele Miragoli, Stefano Rossi, Heimo Ehmke, Julia Gorelik, Wolfgang J. Parak, Neus Feliu, Viacheslav O. Nikolaev

## Abstract

G-protein-coupled receptors (GPCRs) are key mediators of intercellular signalling and targets for the vast majority of current therapeutics. Numerous biochemical and biophysical studies have uncovered that individual GPCRs activated at different subcellular locations can mediate distinct functional effects due to compartmentalization of the downstream second messengers in subcellular nanodomains. However, the reasons why a single receptor subtype activated at the same subcellular compartment such as plasma membrane leads to various functional responses have remained poorly understood. Here, we used β_1_-adrenergic receptor and cardiomyocytes as classical examples of a therapeutically relevant GPCR and highly compartmentalized cell, to study communication between membrane receptors and subcellular cAMP nanodomains based on a combination of nanoscale scanning ion conductance microscopy assisted local receptor activation with Förster resonance energy transfer based live cell imaging of the cAMP pathway. Unexpectedly, this approach revealed that β_1_-adrenoceptors located in cardiomyocyte T-tubular membrane are strongly coupled to the regulation of cardiac contractility, whereas receptors residing in the outer non-tubular membrane preferentially engage nuclear nanodomains linked to hypertrophy. This unique mechanism of spatially segregated signalling was maintained in failing rodent and human hearts, indicating that it is conserved across species and preserved in disease. Finally, we explored potential therapeutic targeting of this mechanism by generating submembrane-specific beta-blockers acting exclusively at the outer non-tubular membrane. These improved compounds could ameliorate hypertrophy and cardiac remodeling without reducing contractility *in vitro* and *in vivo*. Together, these findings uncover an unexpected level of spatial organization of GPCR signaling within the same subcellular compartment and show that not only its identity but also its submembrane location determines cellular signaling and functional response engaged by the same GPCR.

## Main Text

Heart failure is a devastating cardiovascular disease with immense socio-economic costs and extremely high mortality rates, despite the existence of numerous modern therapeutics. Pathological cardiac remodelling which occurs in heart failure, is largely driven by a hyperactive sympathetic nervous system and chronic activation of GPCRs on the surface of heart muscle cells, also known as cardiomyocytes. Classical example of the disease promoting GPCR in this case is the β_1_-adrenergic receptor (β_1_-AR) with its principal downstream second messenger, 3’,5’-cyclic adenosine monophosphate (cAMP), which promotes cardiac remodelling by inducing pro-hypertrophic gene transcription and activating apoptotic signalling pathways. Therefore, pharmacological inhibitors of cardiac β_1_-ARs, commonly known as beta-blockers, are beneficial and indispensable for heart failure therapy (*1*). Despite all benefits, beta-blockers also lead to a marked depression of cardiac contractility, which is due to the importance of the β_1_-AR/cAMP pathway for the regulation of the excitation-contraction coupling in cardiomyocytes (*2, 3*). This contractility impairment underpins reduced physical capacity and poor patient compliance at the beginning of therapy, as well as limitations in the use of beta-blockers at advanced stages of disease or right after myocardial infarction, clinical settings which especially rely on sufficient cardiac performance. The mechanistic basis for the two distinct functional effects of sarcolemmal β_1_-ARs leading to either increased contractility or to cardiac remodelling, is still elusive.

One possible answer might lie in the compartmentation of β_1_-AR-induced cAMP in so-called nanodomains. These are highly regulated subcellular spaces activated by distinct pools of membrane receptors and segregated by a plethora of mechanisms, including cAMP-degrading enzymes called phosphodiesterases (PDEs) (*4, 5*), physical diffusion barriers, intracellular cAMP buffering and liquid phase separation of downstream signalling molecules (*6*). Subcellular compartmentation of local cAMP pools has been shown to explain differential spatial patterns of cAMP and their respective functional responses engaged by β_2_-AR *vs*. β_1_-AR subtypes, whereby β_2_-AR stimulates cAMP production but does not drive cardiac remodelling (*7, 8*). We previously developed a novel combination of nanoscale scanning ion conductance microscopy (SICM) technique and a fluorescent cAMP biosensor based on Förster resonance energy transfer (FRET) to determine the submembrane localization of both functional β-AR subtypes (*8*). We found that β_2_-ARs are located exclusively in the membrane invaginations of cardiomyocytes, called T-tubules, resulting in highly confined β_2_-AR/cAMP signals. In contrast, β_1_-AR can generate more diffuse cAMP signals, originating from both T-tubules and outer membrane regions, referred to as cell crests (*8*). However, it remained unclear if β_1_-ARs at both membrane locations are coupled to distinct functional responses. Here, we aimed to investigate whether β _1_-ARs located in T-tubules *vs*. cell crests might engage distinct subcellular cAMP nanodomains linked to different functional outcomes.

To address this hypothesis, we performed SICM/FRET measurements in adult rat ventricular myocytes expressing a nuclear-targeted cAMP biosensor, Epac1-NLS. Selective stimulation of β_1_-AR using a SICM pipette positioned on cell crests led to significantly stronger nuclear cAMP signals than those triggered by the receptors stimulated in the T-tubules (**Fig. 1a-c**). Likewise, measurements of nuclear cAMP dependent protein kinase (PKA) activity, known to be critically involved in cardiac hypertrophy (*9*), using the AKAR3-NLS biosensor in the same setting showed much more pronounced responses from the crest (**Fig. 1d-f**). This difference was not observed when measuring overall cAMP output and PKA activity in the cytosol of the cells (**Extended Fig. 1**), suggesting similar numbers of activated receptors at both submembrane locations and a high degree of β_1_-AR/cAMP signal compartmentation. This process was at least in part mediated by the action of PDEs, since inhibition of the functionally relevant PDE3 family abolished the difference between responses to T-tubular and crest β_1_-AR stimulation (**Fig. 1g**). Next, we confirmed that distinct coupling of crest *vs*. T-tubular β_1_-AR responses to nuclear PKA activity is maintained in failing rat myocytes isolated from a chronic 16-weeks post-myocardial infarction heart failure model (**Fig. 1h**) and is also present in human failing ventricular myocytes (**Fig. 1i**). PKA activity responses measured in the nucleus upon both T-tubular and crest β_1_-AR stimulation were completely abolished by the hydrophilic beta-blocker sotalol (**Extended Fig. 2a**), suggesting that they do not involve intracellular pools of β_1_-AR (*10, 11*). Likewise, inhibition of the OCT3 transporter, which is responsible for catecholamine import into the cell and potential activation of intracellular β_1_-AR pools, or inhibition of endocytosis using a dynamin inhibitor did not reduce the SICM/FRET signals (**Extended Fig.2b-d**), excluding the involvement of intracellular receptors under our experimental conditions.

**Figure 1.**
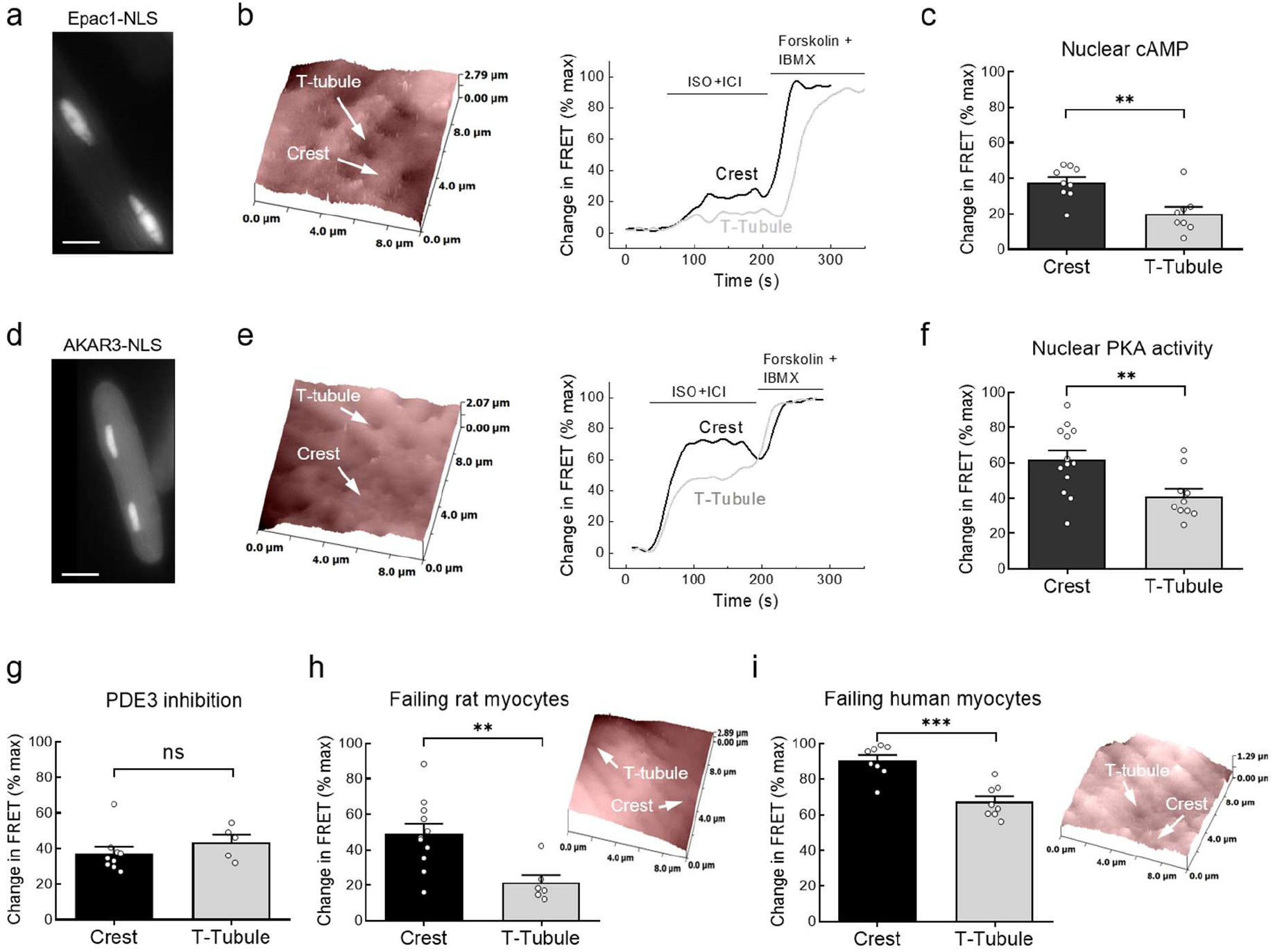
Crest β_1_-ARs are preferentially coupled to nuclear cAMP/PKA signalling. (**a-c**) Rat myocytes expressing nuclear targeted cAMP biosensor Epac1-NLS were topographically scanned and β_1_-ARs were locally stimulated at single T-tubules *vs*. cell crests using SICM pipette filled with isoproterenol (ISO, 50 µM) and the β_2_-AR blocker ICI118,551 (ICI, 5 µM) by applying positive voltage bias to expel femtovolume of negatively charged ISO molecules. Cells were constantly superfused with 50 nM ICI to block β_2_-ARs and ensure local stimulation. At the end of the experiment, bath stimulation with 10 µM forskolin and 100 µM of the pan-PDE inhibitor IBMX was used to evoke maximal sensor response. Crest stimulation induced significantly higher nuclear cAMP levels. n=8-9 cells from 7 rats. (**d-f**) Nuclear PKA activity was measured under the same conditions using the AKAR3-NLS biosensor and showed significantly higher responses to crest β_1_-AR stimulation. n=10-13 cells from 8 rats. (**g**) Pretreatment of rat myocytes with the PDE3 inhibitor cilostamide (10 µM for 5 min prior to SICM/FRET measurements) abolished the difference between T-tubular and crest responses measured as in f (n=5-9 cells from 3-7 rats). Nuclear PKA activity measurements in failing myocytes isolated from rats 16 weeks post myocardial infarction (**h**) and from patients with heart failure (**i**) show maintained significant difference between crest vs. T-tubules responses SCIM/FRET measurements in h and i were performed as in F. n=6-11 cells from 6 failing rats; n=8 cells each from 7 patients. n.s. – not significant, ^**^ p<0.01, ^***^ p<0.001 by mixed ANOVA followed by Wald Chi-squared test.

To assess whether T-tubular β_1_-AR/cAMP signals might be preferentially coupled to the regulation of cardiomyocyte contractility, which could be possible due to the close proximity of L-type calcium channels and sarcoplasmic reticulum calcium release channels (*12*), we stimulated β_1_-AR *via* a SICM pipette at either crests or T-tubules for 1 min and immediately recorded contractility-associated calcium transients. Indeed, T-tubular receptor stimulation led to significantly higher contractile responses as compared to crest (**Fig. 2a-c**), which were maintained even in failing myocytes despite desensitized β_1_-AR responses (**Fig. 2d,e**). These data suggest that the T-tubular pools of β_1_-AR are associated with the regulation of cardiomyocyte contractility, whereas outer membrane receptors located in the crest can trigger stronger nuclear PKA responses associated with cardiac hypertrophy and remodelling. In support of this model, local SICM-based β_1_-AR stimulation at the cardiomyocyte crest led to higher levels of the nuclear early-response transcription factor c-fos, implicated in pathological cardiac growth (*13*) (**Extended Fig. 3**). Our findings are unexpected since the deeper localization of the T-tubular network within the cell would suggest closer proximity to the nucleus and stronger nuclear cAMP responses from crest β_1_-ARs. This scenario was also previously proposed based on experiments with PEGylated isoproterenol, despite its dramatically reduced (>100-fold) affinity for the β_1_-AR (*14*).

**Figure 2.**
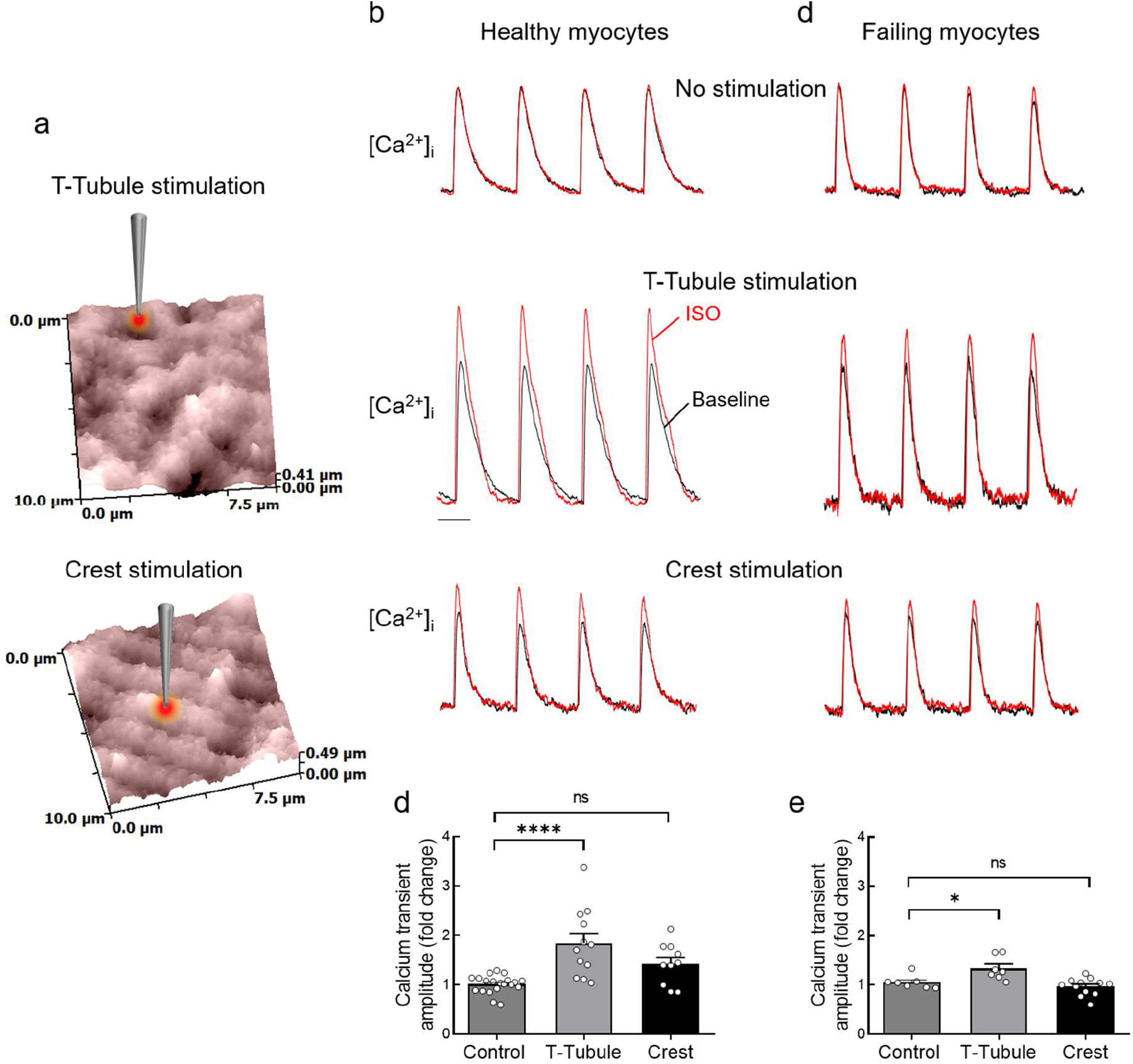
T-tubular β_1_-ARs are preferentially coupled to increased cell contractility. Rat myocytes were loaded with the calcium indicator Fluo4, scanned (**a**) and electrically paced to record basal Ca^2+^ transients (**b**, black traces). Next, cells were stimulated for 1 min with ISO in the presence of ICI applied either into T-tubules or on the crests, as described in Fig. 1, to record stimulated Ca^2+^ transients (**b**, red traces). (**c**) Analysis of Ca^2+^ transient amplitudes from experiments from A-B shows significantly higher responses to T-tubular β_1_-AR stimulation. n=10-20 cells from 3 rats. (**d**) This difference was maintained in failing myocytes isolated from rats after myocardial infarction, despite apparent β_1_-AR desensitization. n=7-13 cells from 3 failing rats. n.s. – not significant, ^*^ p<0.05, ^****^ p<0.0001 by mixed ANOVA followed by Wald Chi-squared test.

To leverage this distinct β_1_-AR nanodomain compartmentation for heart failure therapy, we sought to develop a beta-blocker capable of inhibiting this receptor only at the cell crest. Since cardiac T-tubules serve as a size exclusion filter of around 10-12 nm (*15*), we first used 18 nm gold nanoparticles and covalently linked them with the beta-blocker atenolol *via* a flexible linker attached to an organic polymer (**Fig. 3a**), resulting in 18 nm atenolol@Au NPs. We confirmed their beta-blocker activity and receptor binding capacity in a β_1_-AR expressing cell line (**Extended Fig. 4**). Incubation of adult rat ventricular myocytes with atenolol@Au NPs prevented hypertrophy induced *in vitro* by 24 hour β-adrenoceptor agonist isoproterenol (ISO) treatment, comparable to the effect of regular atenolol (**Fig. 3b**). However, atenolol@Au NPs, as opposed to regular atenolol, had no effect on cardiomyocyte contractility measured by sarcomere shortening *in vitro* (**Fig. 3c**). To measure the response in intact hearts, we used a classical mouse Langendorff preparation (**Fig. 3d**) and treated these hearts with atenolol@BSA@Au NPs having 5 nm core Au NP and albumin molecules between the linker and the NP core (**Fig. 3a**, resulting in a hydrodynamic diameter of >14 nm). Utilizing bovine serum albumin (BSA) enables these nanoparticles to cross capillary barriers and reach the heart muscle tissue (**Extended Fig. 5**). These nanoparticles abolished ISO-induced upregulation of c-fos as an early hypertrophy response gene (**Fig. 3e**) but, as opposed to regular atenolol, did not diminish cardiac contractility in the Langendorff system (**Fig. 3f**) or *in vivo* upon infusion into mice (**Extended Fig. 6**).

**Figure 3.**
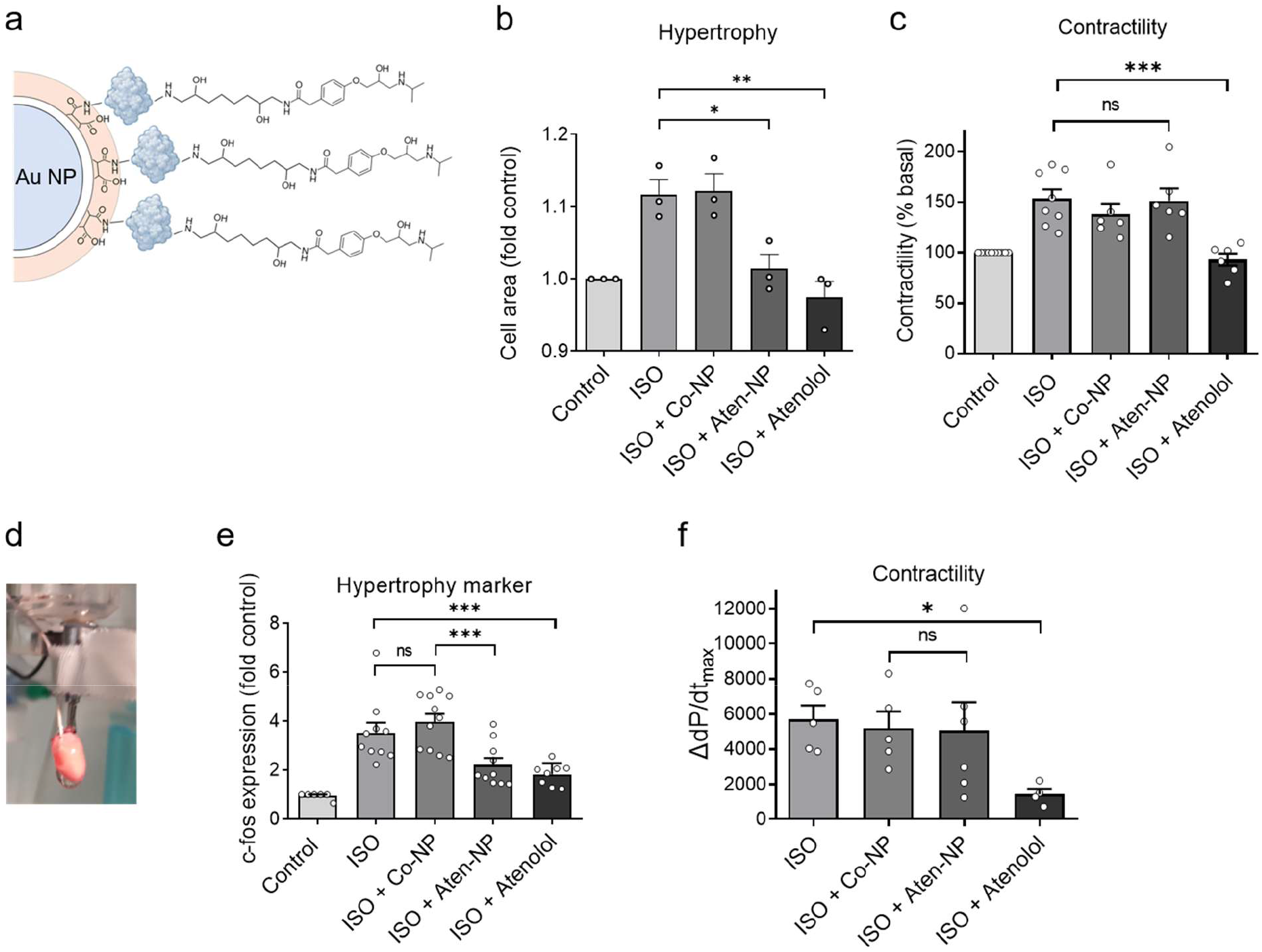
Selective inhibition of crest β_1_-AR prevents myocyte hypertrophy without contractility decline *in vitro*. **(a)** Schematic structure of gold nanoparticles (NP) coated with an organic polymer covalently linked to atenolol (atenolol@Au NPs with 18 nm gold NP cores or alternatively with atenolol-modified albumin (atenolol@BSA@Au NPs with 5 nm NP cores and hydrodynamic size of >14 nm, albumin is schematically shown as blue-gray globular structure). (**b**) Hypertrophy measurement in adult rat myocytes treated for 24 h with 10 nM ISO together with or without atenolol@Au NPs (Aten-NP), Au NPs (Co-NP, with organic polymer but without atenolol, both used at 100 nM; referring the NP core concentration) or pure atenolol (100 nM, referring to the atenolol concentration; NP and atenolol concentrations cannot be directly compared). Data are from n=3 independent isolations with at least 100 cells counted per heart. (**c**) Contractility measurements in cells acutely treated with the same compounds (n=6-7 cells each). (**d**) Langendorff mouse hearts were perfused for 40 min with vehicle (Control), 10 nM ISO alone or together with control BSA@Au NPs (Co-NP) or atenolol@BSA@Au NPs (Aten-NP), both at 100 nM, or regular atenolol (500 nM) to measure c-fos early response hypertrophy gene expression (**e**, n=6-11 hearts) and contractility expressed as maximal rate of pressure change over time (ΔdP/dt_max_) at peak ISO response (**f**, n=4-6 hearts). n.s. – not significant, ^*^ p<0.05, ^**^ p<0.01, ^***^ p<0.001 by one-way ANOVA followed by Sidak multiple comparison test.

To analyze the effect of atenolol nanoparticles on heart failure development *in vivo*, we induced myocardial infarction (MI) in mice and injected them with similar atenolol nanoparticles containing mouse albumin (atenolol@MSA@Au NPs) once weekly starting from 14 days and until 8 weeks after MI. Surprisingly, there was no effect of this treatment on heart-to-body weight ratio and on cardiac performance in mice (**Fig. 4a,b**) which could be due to poor pharmacokinetic properties of systemically administered gold nanoparticles, leading to predominant delivery to liver and spleen (*16*). To enable better *in vivo* pharmacokinetics, we next generated antibody conjugates containing unspecific immunoglobulin G (IgG) antibodies covalently linked to atenolol *via* mouse albumin molecules and having >12 nm size (atenolol@MSA@IgG, see **Supplementary Material**), also called Atenolol-IgG for simplicity. Atenolol-IgG effectively blocked β_1_-AR in a receptor expressing cell line and *in vitro*, showing binding affinities comparable to regular atenolol (**Extended Fig. 7a-f**), and retained selectivity for β _1_-AR, not affecting β _2_-AR/cAMP signalling (**Extended Fig. 8**). When we incubated adult mouse cardiomyocytes with Atenolol-IgG prior to SICM/FRET experiments, it completely blocked cytosolic cAMP responses evoked by β _1_-AR located at the crest but not at T-tubules of the cells (**Extended Fig. 7g-i**). Similar to atenolol nanoparticles, they also inhibited ISO-induced hypertrophy and did not affect contractility of isolated myocytes (**Extended Fig. 9a**,**b**). This correlated well with the effect of Atenolol-IgG on ISO induced phosphorylation of PKA substrates involved in nuclear (pCREB) *vs*. contractile responses (**Extended Fig. 10**). Analysis of Atenolol-IgG content in mouse plasma at various time points after injection revealed a favorable pharmacokinetic profile with half-life of ~7 days (**Extended Fig. 9c**). *In vivo*, Atenolol-IgG could also prevent pathological cardiac growth and contractility decline after MI (**Fig. 4c,d**). Finally, we used a highly standardized pressure overload induced heart failure model in mice following transverse aortic constriction (TAC) and treated these mice once weekly with Atenolol-IgG *vs*. Control-IgG starting at 14 days and until 8 weeks after TAC. Here, Atenolol-IgG effectively ameliorated TAC induced increase in heart weight, cardiomyocyte hypertrophy, without reduction of cardiac contractility (**Fig. 4e-h, Extended Fig. 11**).

**Figure 4.**
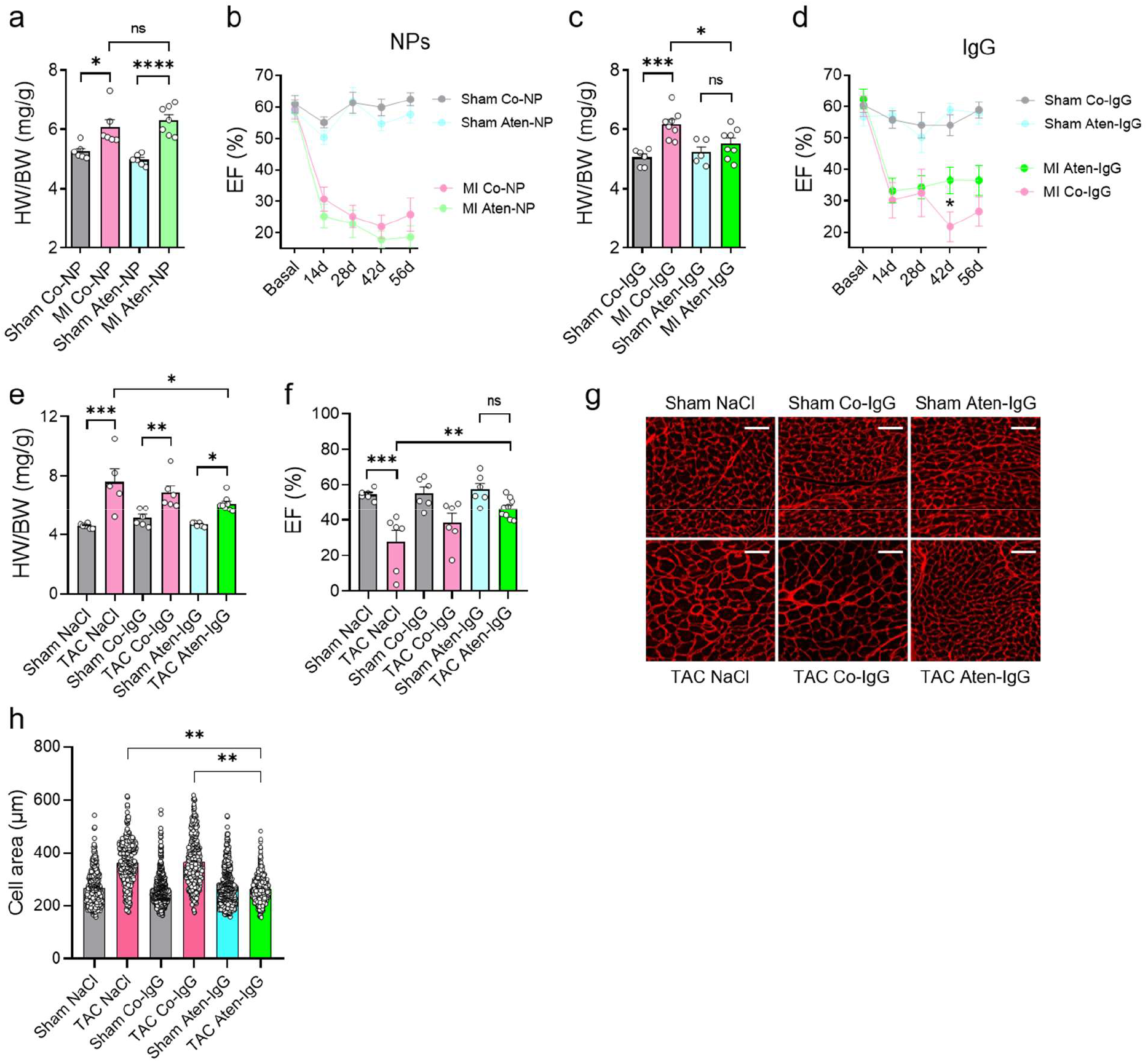
Atenolol-linked antibodies prevent cardiac hypertrophy without reducing cardiac contractility *in vivo*. Mice were subjected to myocardial infarction (MI, **a-d**) or transverse aortic constriction (TAC, **e-f**) surgeries and injected once weekly intravenously (2.4 mg/kg) with MSA@Au NP (Co-NP) vs. atenolol@MSA@Au NP (Aten-NP, **a-b**), or subcutaneously (1 mg/kg) with MSA@IgG (Co-IgG) or atenolol@MSA@IgG (Aten-IgG, **c-f**) starting 2 weeks after surgery. Cardiac hypertrophy was assessed based on heart by body weight ratios (HW/BW) (**a**,**c**,**e**), and contractile performance was compared by measuring ejection fraction of the heart by serial echocardiography performed every two weeks after MI (**b**,**d**) or 8 weeks after TAC (**f**) surgeries. n=6-9 mice each. (**g**,**h**) Analysis of cardiomyocyte cell area from sham and TAC heart sections 8 weeks after surgery. Scale bars, 50 µm. Data are from at least 100 cells counted per heart from n=5-6 hearts per group. n.s. – not significant, ^*^ p<0.05, ^**^ p<0.01, ^***^ p<0.001, ^****^ p<0.0001 by one-way ANOVA followed by Sidak multiple comparison test in A,C,E,F, and ^**^ p<0.01 by mixed ANOVA followed by Wald Chi-squared test in H.

In summary, we demonstrate that β_1_-AR pools located in the T-tubular membrane nanodomains are preferentially coupled to cardiac contractility whereas those located on the cell crest preferentially regulate hypertrophy. This provides the evidence that the same GPCR depending on its “geographic location” within one subcellular compartment, in this case sarcolemmal membrane, can be functionally segregated to trigger distinct downstream signalling pathways and physiological effects. This paradigm can be leveraged for the development of a new generation of nanodomain-specific beta-blockers based on size exclusion of nanoparticles or antibodies covalently decorated with atenolol molecules which are capable of inhibiting only β_1_-ARs located at the cardiomyocyte outer membrane. These improved beta-blockers can effectively prevent cardiac hypertrophy and heart failure progression without reducing contractility, thereby allowing more precise treatment for heart failure patients. We believe that this principle can be potentially transferred to other GPCRs and disease states.

## Acknowledgments

We thank Viktoria Hänel for excellent technical assistance. We would also like to thank the Facility for Imaging by Light Microscopy and the Centre of Excellence Cellular Mechanosensing and Functional Microscopy at Imperial College London.

## Author contributions

Investigation: AF, BQ, MF, GB, KM, SK, XS, HS, JLS-A, RM, PTW, MG, DR, CEM, NIB, SR, MM and VON Conceptualization and Supervision: HE, JG, WJP, NF, VON Writing – original draft: VON Writing – review & editing: all authors

## Data availability

All data are available in the main text or the supplementary materials.

## Supplementary Materials

Materials and Methods Table S1

Extended Figs. 1 to 11

References (*17*–*27*)

