## Supplemental Methods and Figures for "Submembrane geography dictates GPCR signalling compartmentation and physiological function"

**The PDF file includes:**

Materials and Methods
Extended Figs. 1 to 11
Table S1

### Materials and Methods

Rat cardiomyocyte isolation

Adult rat ventricular myocytes were isolated as previously described (*8*). Failing rat myocytes were isolated from rats 16 weeks after myocardial infarction from the previously described and characterized cohort of rats (*17*). Freshly isolated cells were plated on laminin collated glass bottomed dishes and transduced with adenoviruses encoding indicated fluorescent biosensors and used for microscopy experiments 40-48 h after transduction. Hypertrophy assay was performed with freshly isolated cells by incubating them for 24 h in presence of 10 µM 2,3-butanedione monoxime and 10 nM isoproterenol to stimulate cell growth (*18*) combined with beta-blocker preparations as indicated in figure legends. Cells were then fixed, photographed using Leica DMi1 microscope equipped with MC120 HD camera, and cell surface area was measured using ImageJ software after manual selection of individual single cells and application of the Wand tool for automated area outlining and measurement.

Human cardiomyocyte isolation

Human ventricular tissue samples were obtained from heart failure patients during open heart surgery. For all ventricular tissue samples, patients provided written informed consent. The study followed the Declaration of Helsinki and was approved by the Ethical Committee of the Aerztekammer Hamburg (PV7015). Clinical characteristics of patients are described in the **Table S1**. Fresh tissue samples were used for cardiomyocyte isolation as previously described (*19*). After isolation, cells were resuspended in human myocyte culture medium (minimum essential medium containing creatine (12 mM), taurine (20 mM), penicillin/streptomycin (1 %), L-glutamine (1 %), BSA (0.1 %), blebbistatin (12 µM), and fetal calf serum (FCS, 2.5 %), and plated on laminin-coated dishes. 2 h later, FCS-free culture medium was added and adenoviral transduction was performed as for rat myocytes described above.

Scanning ion conductance microscopy (SICM) and Förster resonance energy transfer (FRET)

SICM is a scanning probe microscopy technique based on measuring ion current flowing through a tip of a glass nanopipette (~100 MΩ resistance, corresponding to 40-50 nm inner diameter which determines the spatial resolution of the scan). The pipettes were filled with buffer A (144 mM NaCl, 5.4 mM KCl, 1 mM $MgCl_2$, 1 mM $CaCl_2$ and 10 mM HEPES, pH=7.3) containing isoproterenol and other receptor ligands as indicated. The lower vertical (z axis) position of the pipette tip was kept constant relative to the cell surface to perform 10x10 µm cell surface scan at -200 mV pipette voltage (*20*). In this study, we used the hopping probe ion conductance microscopy (*21*) and the ICnanoS scanning system (Ionscope, Melbourn, UK) or similar custom designed SICM system (ICAPPIC, London, UK) positioned on the Nikon Eclipse T*i* inverted fluorescence microscope. During application of receptor ligands, we minimized the hopping amplitude and superfused the cell with the buffer A to ensure a strictly local receptor stimulation (<500 nm) as previously described (*8*). Local nanoscale receptor stimulation was achieved by applying a positive voltage bias (-500 mV) to the pipette to expel femtoliter amounts of negatively charged isoproterenol with comparable final concentrations of the ligand at T-tubular and Crest membranes as described (*22*).

FRET imaging of cAMP or PKA activity in stimulated cells was performed using a custom build imaging system, consisting of a 440 nm pE-1000 light emitting diode (CoolLED) to excite the biosensor, the microscope (as above) and a DV2 DualView beam splitter (Photometrics) equipped with 505dcxr dichroic mirror, D480/30 and D535/40 emission filters. Fluorescence in individual channels was recorded by the ORCA-03G camera (Hamamatsu) using the open source MicroManager 1.4 software (*23*). FRET ratios were corrected for the bleedthrough of the CFP into the YFP channel and analysed using Origin 8.0 software as described (*24,25*).

Contractile calcium measurements following local receptor stimulation were performed in Fluo-4-loaded cardiomyocytes paced at 0.5 Hz. Cells were superfused with buffer A containing 1 mM $Ca^{2+}$ and exposed to ICI 118,551 (50 nM) and blebbistatin (5 µM) in the bath. For local stimulation, isoproterenol (ISO, 30 µM) together with ICI 118,551 (5 µM) was loaded into the pipette. After acquisition of the SICM scan, the pipette was positioned approximately 500 nm above the selected membrane site, corresponding to either a T-tubule opening or a crest region, and ISO was applied locally for 1 min using a positive pulse of 500 mV. Calcium transients were recorded before ISO application and at 1, 2, and 5 min after stimulation. Experiments were performed at room temperature.

Langendorff perfused mouse heart model

Carefully dissected hearts were placed in ice-cold modified Krebs-Henseleit solution (mM: NaCl 119, $NaHCO_3$ 25, KCl 4.6, $KH_2PO_4$, 1.2, $MgSO_4$ 1.1, $CaCl_2$ 2.5, glucose 8.3 and Na-pyruvate 2; pH 7.4, 95 % $O_2$/5 % $CO_2$). The aorta was cannulated and quickly attached to the perfusion system (Hugo Sachs Elektronik/Harvard Aparatus, Germany) for retrograde perfusion with the same buffer at constant pressure (80 mm Hg) as described (*26*). Octopolar catheter was placed into left ventricle to record contractility and heart rate. To measure developed pressure, a Millar catheter was additionally inserted into left ventricle. Hearts that did not regain a pink coloration or a spontaneous heartbeat after 20 min equilibration were discarded. After equilibration, hearts were infused with beta-blocker preparations for 5 min followed by beta-blocker together with isoproterenol for 40 min, after which they were frozen for gene expression analysis.

Quantitative real time polymerase chain reaction

Total RNA from Langendorff perfused mouse hearts or frozen hearts from in vivo experiments was isolated using the RNeasy fibrous tissue mini kit (Qiagen). mRNA was then transcribed into cDNA using the iScript cDNA synthesis kit (BioRad) with 200 ng of RNA per reaction. Rotor-Gene SYBR Green PCR Kit (Qiagen) was used for quantitative real-time PCR reaction (qRT PCR) in a Rotor Gene Q cycler (Qiagen). Samples were run in triplicates using the following conditions: 10 min at 95°C, followed by 40 x three step cycling (10 s at 95°C, 15 s at 60°C, 20 s at 72°C), and running a melting curve (72 – 95°C, 0.5°C steps). The following primer pairs were used: Gapdh (forward: 5’-CGTCCCGTAGACAAAATGGT -3’; reverse: 5’-GAATTTGCCGTGAGTGGAGT -3’), c-fos (forward: 5’- CCACCGACCTGCCTGCAAGA -3’; reverse: 5’- TGGCTTGGGCTCAGGGTCGT -3’).

Mouse surgeries

All animal procedures were approved by the local animal welfare authority BJV Hamburg (approval number N080/2022) and conformed to the guidelines from Directive 2010/63/EU of the European Parliament on the protection of animals use for scientific purposes.

*In vivo hemodynamic* – Left ventricular (LV) function was assessed using a 1.2-F microconductance catheter introduced into the LV via the carotid artery under isoflurane anesthesia (1.5–2 %) and buprenorphine (0.1 mg/kg i.p.). Pressure-volume loops were recorded for 30–45 minutes using a rectal thermometer-controlled heating plate. For therapeutic studies, nanoparticle-based beta-blockers were infused via the left jugular vein (0.26 µl/g/min for 10 min). Volume calibration was performed via wash-in technique with 10% NaCl. Measurements conducted during brief apnea.

*Myocardial infarction (MI)* – After randomization, 11- to 13-week-old mice (both males and females) were initially anaesthetized with 4% isoflurane in 100 % oxygen and maintained via endotracheal intubation (1.5–2.5 %) with mechanical ventilation. Pre-surgical analgesia consisted of buprenorphine (0.1 mg/kg s.c.) administered 30 minutes prior to induction. Following a left lateral thoracotomy at the third intercostal space, the left anterior descending artery (LAD) was identified, after opening the pericardium, and ligated with a non-resorbable suture approximately 2 mm below the left auricle. Successful induction of ischemia was confirmed by visible myocardial blanching distal to the ligation. Postoperative care included carprofen and buprenorphine (5 mg/kg s.c. for, 0.1 mg/kg s.c.) injection, metamizole via drinking water, and monitored recovery on a heating pad. Sham-operated animals underwent the identical surgical preparation and thoracotomy, except for the ligation of the LAD. 8 weeks after surgery, heart structure and contractile function were analyzed by echocardiography, performed using the Vevo 2100 system (VisualSonics) equipped with a 25-55-MHz transducer (MX-550D) as described (*27*).

*Transverse aortic constriction (TAC)* - After randomization, 9- to 12-week-old mice (both males and females) were anaesthetized with a combination of medetomidine (0.5 mg/kg body weight), midazolam (5 mg/kg body weight), and fentanyl (0.05 mg/kg body weight), injected once intraperitoneally, and subjected to a suprasternal incision. The aortic arch was exposed in a minimal invasive manner and constricted between the first and second trunk, using a 27 G spacer and 6–0 polyviolene suture. Anesthesia was antagonized with intraperitoneal injection of atipamezole (2.5 mg/kg body weight) and flumazenil (0.1 mg/kg body weight). Operated mice received analgesic therapy with buprenorphine (0.05 mg/kg subcutaneously, 30 min prior to and every 8 h for 3 days after surgery) and carprofen (5 mg/kg subcutaneously, 30 min prior to and every 24 h for 3 days after surgery), as well as oral metamizol therapy for 1 week after surgery. 3-5 days post-surgery, pressure gradients across the transverse aortic constriction (TAC) region were measured using transthoracic echocardiography utilizing a 20 MHz Doppler velocity probe (Visual Sonics Vevo 3100, Toronto, Canada). When the pressure gradient was above 50 mmHg, the constriction was considered sufficiently good and mice were used in experiments.

Immunoblot analysis

Adult rat vertricular myocytes were pre-incubated with MSA@IgG (Control IgG) ot atenolol@MSA@IgG (Atenolol IgG), both at 200 µg/mL, for 30 min before stimulating with 10 nM isoproterenol. 10 minutes after stimulation, cells were immediately lysed with RIPA buffer (20 mM Tris pH 7.4, 150 mM NaCl, 1 % Triton X-100, 0.5 % Sodium deoxycholate, and 0.1 % SDS) supplemented with cOmplete protease inhibitor-cocktail and PhosSTOP (both from Merck, Darmstadt, Germany). Equal amount of proteins was loaded onto the gel, transferred to a nitrocellulose membrane and proteins were detected using antibodies specific for phospho-CREB Ser133 (catalog number 9198; Cell Signaling Technology, Leiden, NL), phospho-PLN S16 (A010-12AP; Badrilla, Leeds, UK), total PLN (126174; Abcam, Cambridge, UK) and GAPDH (5G4; HyTest Ltd, Turku, Finland). Subsequently, HRP-labelled anti-mouse or anti-rabbit antibodies (both from Bio-rad, Dreieich, Germany) were used. The blots were developed using SuperSignal™ West Pico PLUS chemiluminescense substrate (Thermo Scientific, Dreieich, Germany). The signals were quantified using ImageJ.

Cell size analysis

Biventricular paraffin cross-sections (3 µm) were deparaffinized in xylene and rehydrated through a graded ethanol series. For antigen retrieval, sections were heated in sodium citrate buffer (10 mM sodium citrate in $ddH_2O$, 0.05 % Tween 20, pH 6.0) for 15 minutes at 200 W using a microwave. Following retrieval, membranes were stained Alexa Flour 555 labeled Wheat Germ Agglutinin (WGA, 75 µg/mL, Sigma-Aldrich) for 30 minutes. To quench tissue autofluorescence, sections were incubated in 0.2 % sudan black diluted in 70 % ethanol for 30 min. Finally, slides were mounted using Fluoromount-G (Invitrogen) and imaged using a Pannoramic MIDI II slide scanner (Sysmex) equipped with a 20x objective (Plan-Achromat, Zeiss). For morphometric analysis, regions of interest (ROIs) containing strictly cross-sectionally oriented cardiomyocytes were selected along the inner circumference of the left ventricle. After image normalization, a machine-learning-based pixel classifier (Ilastik, version 1.4.0.post1) was trained on a subset of images to generate probability maps. These maps were then processed in ImageJ (version 1.48g4) using an automated thresholding to identify individual cardiomyocytes. Particle analysis was performed using a size filter of 1500-6000 pixel^2 and a circularity index of 1.5-1.0 to ensure accurate crosssectional-area quantification. 5 mice were used per group, with 150-400 cells measured per mouse and 600-1000 total cells per group.

Single-cell contractility measurements

Adult rat vertricular myocytes were plated onto glass bottomed dished and preincubated with NP or IgG preparations as described in figure legends. Contractility measurement were performed using multicell CytoCypher system (IonOptix) at 1 Hz pacing frequency at basal conditions and under ISO stimulation as previously described (*27*).

Statistics

Echocardiographic, morphometric, SICM/FRET data were analyzed using the Origin Pro 8.5 (OriginLab Corporation, Northhampton, MA), GraphPad Prism 7 (GraphPad, San Diego, CA) and R 3.6.3 software (R Foundation for Statistical Computing, Vienna, Austria). Normal distribution was tested by Kolmogorov-Smirnov test or by Shapiro-Wilk normality test, the latter was used only for groups of data with $n<6$. Differences between the groups of normally distributed data were analyzed using mixed ANOVA followed by Wald Chi-squared test, paired or unpaired t-test for simple two-group comparison or one-way ANOVA followed by Sidak test for multiple groups. Groups of unpaired non-parametric data were analyzed by Mann-Whitney test, whereas Friedman test was used for paired non-parametric data. For multiple comparison, parametric Sidak's multiple comparisons test for preselected groups, Tukey's test for paired data or nonparametric Dunn's test for data with skewed distribution were used, as appropriate. Sample sizes were 3-9 mice/rats/patients and 6-17 measured cells. No sample exclusion was performed. Representative images were selected to represent the means of the quantified data.

### Extended Figures

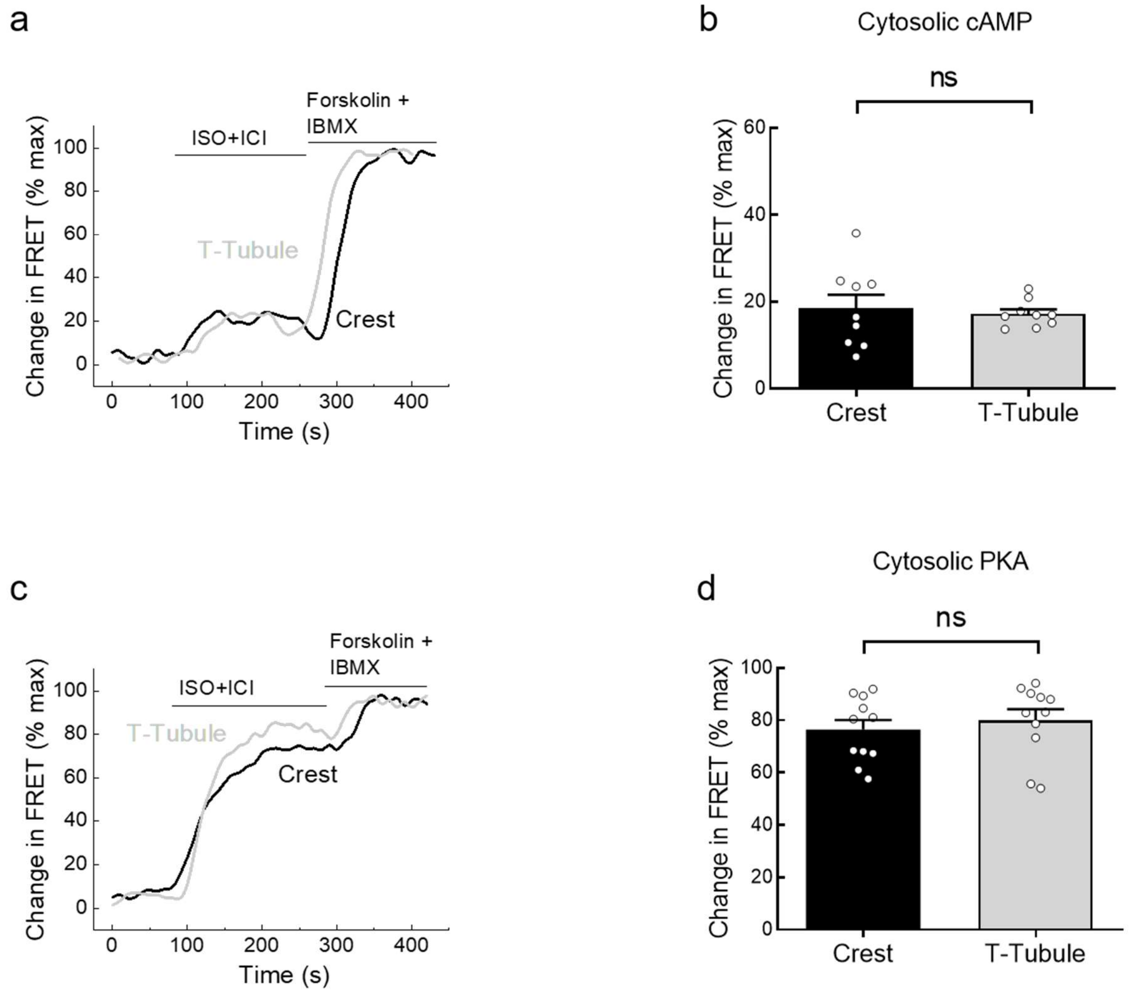


**Extended Figure 1. Cytosolic cAMP and PKA responses after local T-tubule and crest stimulation of β$_1$-AR.** Rat ventricular myocytes expressing the cytosolic Epac1-camps or AKAR3-NES biosensors were locally stimulated at single T-tubules or cell crests using an SICM pipette containing isoproterenol (ISO, 50 µM) and the β$_2$-AR blocker ICI118,551 (ICI, 5 µM) as described in Fig. 1. Cells were constantly superfused with 50 nM ICI to block β$_2$-ARs and ensure local stimulation. Maximal sensor responses were obtained by bath application of forskolin (10 µM) and IBMX (100 µM). (**a**) Representative FRET traces showing cytosolic cAMP signal output measured with the cytosolic Epac1-camps biosensor. (**b**) Quantification of the c cAMP-FRET respones upon stimulation of β$_1$ARs in crest and in the T-tubules of rat cardiomyocytes. n=9 cells from 3 rats each. n.s. – not significant by mixed ANOVA followed by Wald Chi-squared test. (**c**) Representative FRET traces showing cytosolic PKA activity measured with the cytosolic AKAR3-NES biosensor. (**d**) Quantification of the changes in FRET ratios upon stimulation of β$_1$ARs in crest and in the T-tubules of rat cardiomyocytes. n=11 cells from 6 rats. n.s. – not significant by mixed ANOVA followed by Wald Chi-squared test.

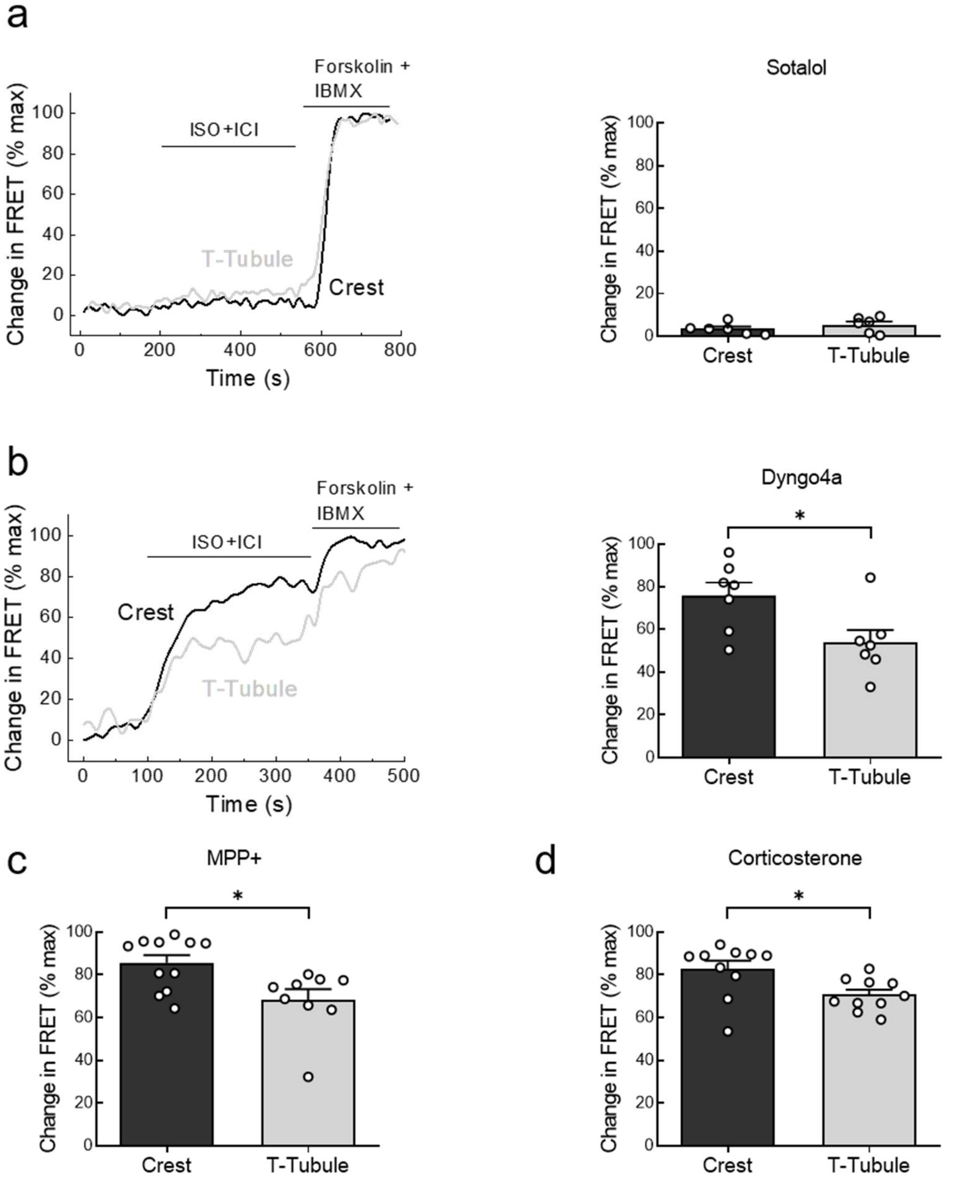


**Extended Figure 2. Nuclear PKA responses to local β1-AR stimulation are mediated by cell-surface receptors and do not involve intracellular receptor pools.** PKA activity in the nucleus was measured using AKAR3-NLS biosensor after local β1-AR stimulation by SICM pipette (as described in Figure 1) at T-tubules and cell crests. The nuclear responses were completely abolished by the hydrophilic β-blocker (**a**) sotalol (100 μM), indicating that they do not depend on intracellular β1-AR pools. Inhibition of endocytosis with a dynamin inhibitor (**b**, Dyngo4A, 30 μM, 30 min pretreatment) did not reduce the SICM/FRET responses, likewise inhibition of the OCT3 transporter (**c,d**) using 100 μM MPP+ (30 min pretreatment) or 20 μM corticosterone (20 min pretreatment). n=6-11 cells each from 4-6 rats. * $p<0.05$ by mixed ANOVA followed by Wald Chi-squared test.

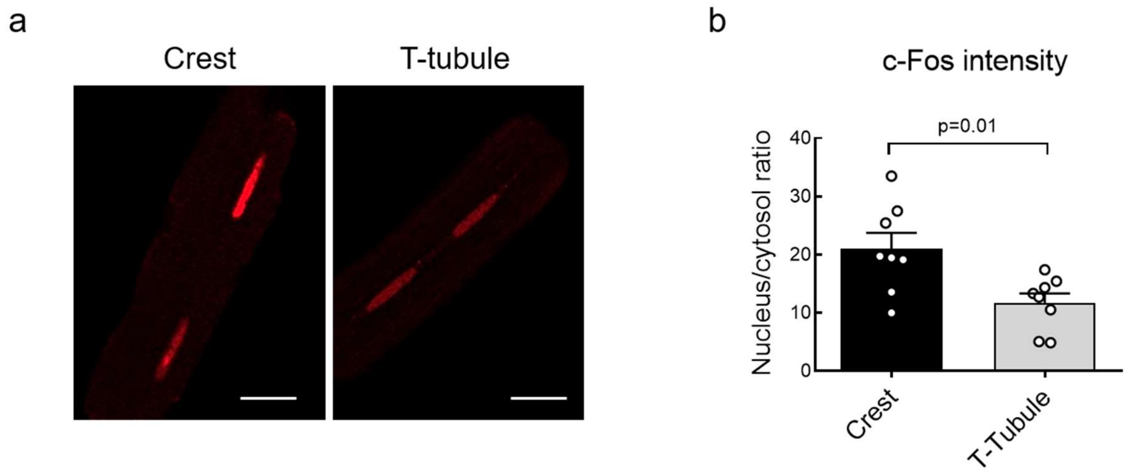


**Extended Figure 3. Crest-localized $\beta_1$-AR stimulation increases nuclear c-Fos expression.** Rat cardiomyocytes were locally stimulated for 45 min *via* SICM pipette as described in Fig. 1 either at the crest or at the T-tubule, fixed and stained for nuclear expression of the early-response transcription factor c-Fos, which is associated with pathological cardiac growth and remodeling. (**a**) Representative confocal images. (**b**) Quantification of nuclear c-Fos expression as the nucleus/cytosol intensity ratio, showing enhanced nuclear accumulation after crest stimulation. n=8 cells each from 4 rats. ** $p<0.01$ by t-test.

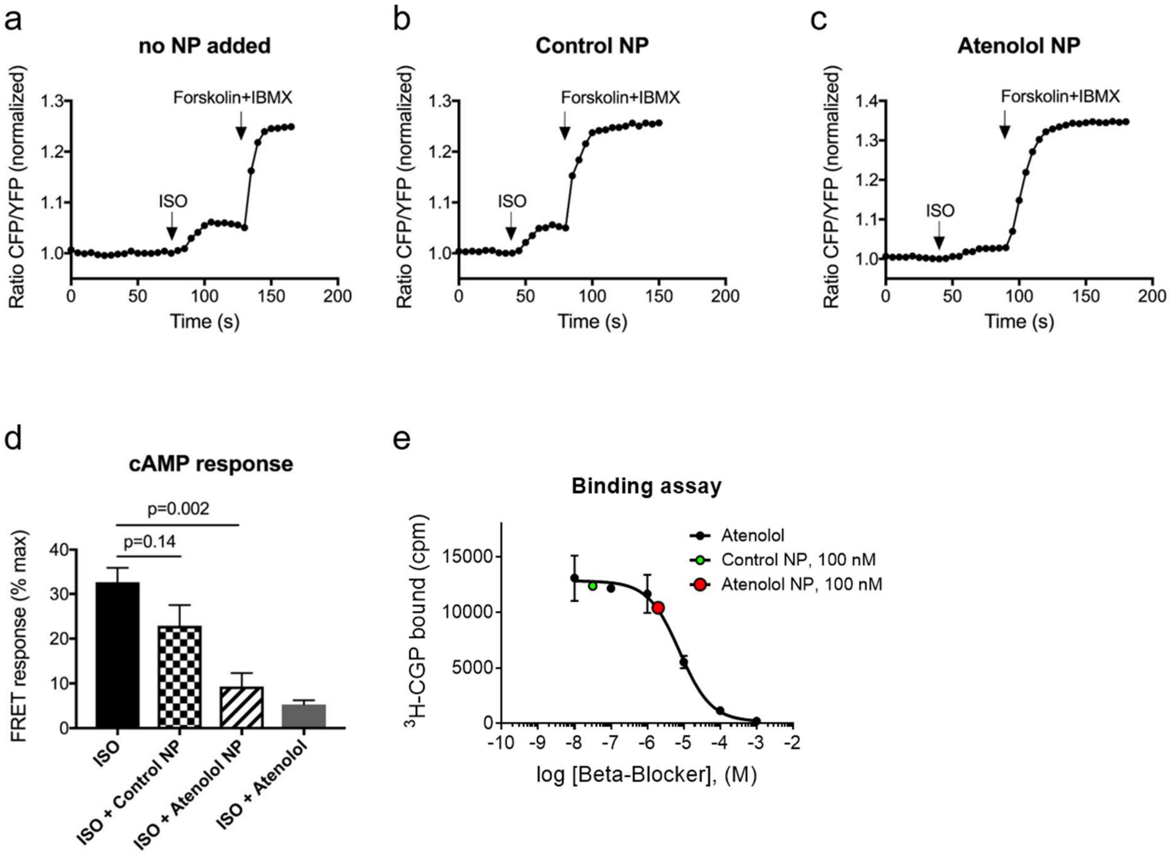


**Extended Figure 4. Beta-blocker activity and receptor binding capacity of Atenolol NP.** (**a-c**) Representative FRET traces showing cAMP responses to isoproterenol (ISO, 0.03 nM) in HEK293 cells expressing $\beta_1$-AR together with the cytosolic cAMP biosensor Epac1-camps recorded after 30 min pretreatment with vehicle (no NP added), Au NPs (Control NPs, 100 nM) or atenolol@Au NPs (Atenolol NPs, 100 nM). Maximal sensor response was obtained by stimulation with forskolin (10 µM) and IBMX (100 µM). (**d**) Quantification of FRET responses shown in A-C along with the ISO response in presence of 30 nM of regular atenolol. n=8-10 cells per condition. ** $p<0.01$, ns – not significant by one-way ANOVA followed by Sidak multiple comparison test. (**e**) Radioligand binding assay performed on membranes from the HEK293 cells expressing $\beta_1$-AR incubated with the radioligand $^3$H-CGP12177 and increasing concentrations of unlabeled atenolol in presence of Au NPs (Control NP) or atenolol@ Au NPs (Atenolol NP), both at 100 nM. We could not use higher, increasing concentrations of Au NPs because their colloid stock solutions tend to aggregate at higher concentrations (>300 nM, used to prepare 1:3 dilutions for the assay). Atenolol NP but not control NP bind to the receptor and displace the unlabeled atenolol. $IC_{50}$ value for atenolol was 7.7±0.2 µM. Data are mean±SE from 4 independent assays. Red data point corresponds to 100 nM Atenolol NP and 2.3 µM atenolol, suggesting that each Au NP contains at least ~23 fully active atenolol molecules when calculated using the molecular weight of the Au NP core only ($8.08\cdot10^5$ g/mol).

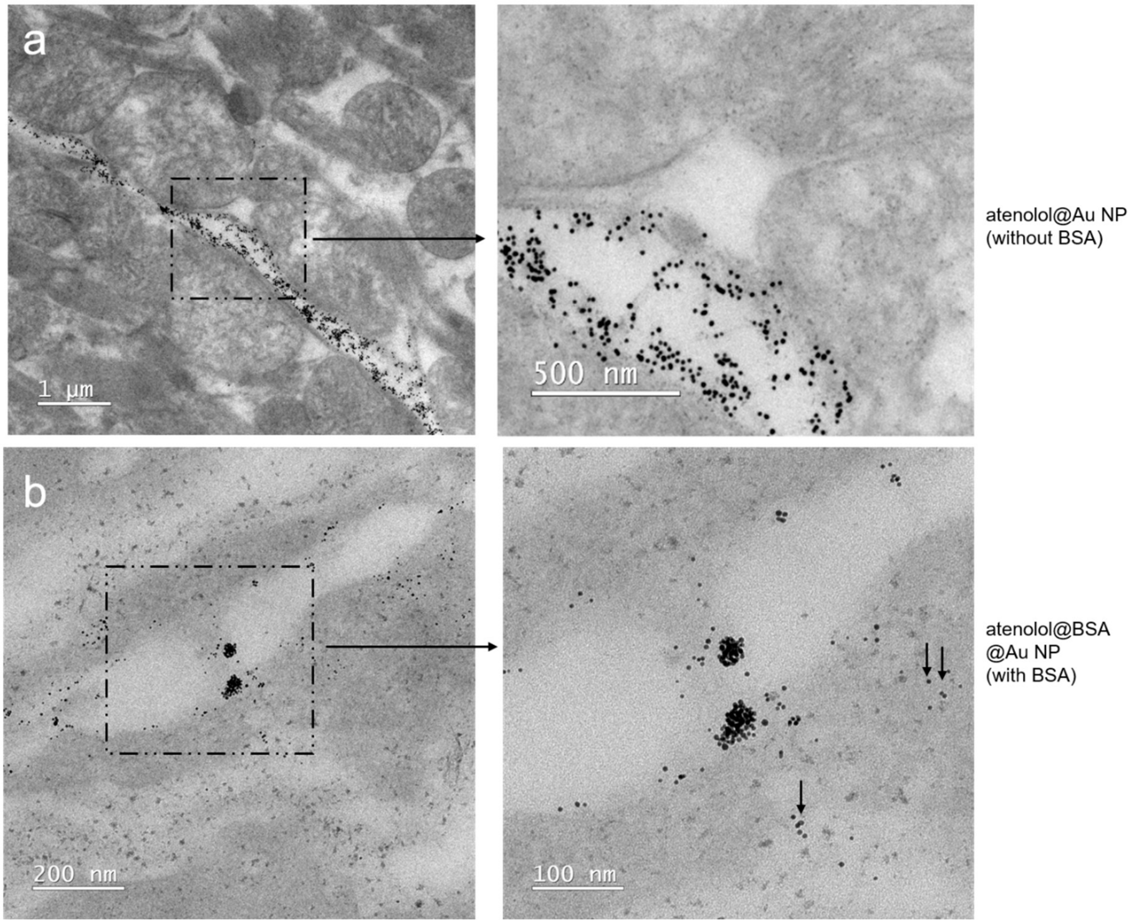


**Extended Figure 5. Nanoparticle distribution in Langendorff-perfused mouse hearts.** Representative transmission electron microscopy (TEM) images show that atenolol@Au NPs (18 nm core diameter) accumulate within capillaries and remain largely confined to the vascular space **(a)**, whereas atenolol@BSA@Au NPs (5 nm core diameter, but due to the BSA conjugation having a similar hydrodynamic diameter as the 18 nm core diameter NPs) cross capillary barriers and reach the surrounding myocardium **(b)**. Representative images, n=8-10.

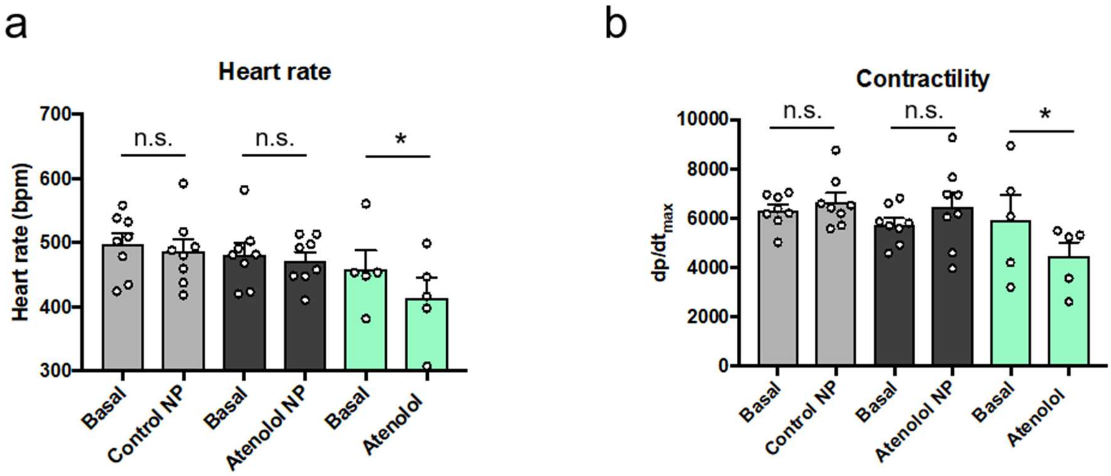


**Extended Figure 6. *In vivo* hemodynamic responses to MSA@Au NPs (Control NP) or atenolol@MSA@Au (Atenolol NP) and regular atenolol**. Control NP, Atenolol NP or regular atenolol were infused into anesthetized mice *via* jugular vein (100 ng/g/min for NPs; mass concentration referring to the NP core concentration determined by UV/vis absorption spectroscopy, and 200 ng/g/min for regular atenolol; mass concentration referring to atenolol). Heart rate (**a**) and cardiac contractility (**b**) were measured before and 10 min after infusion. Atenolol NP did not affect contractility as opposed to regular atenolol. n=5-8 mice per group, n.s. – not significant, * $p<0.05$ by paired t-test.

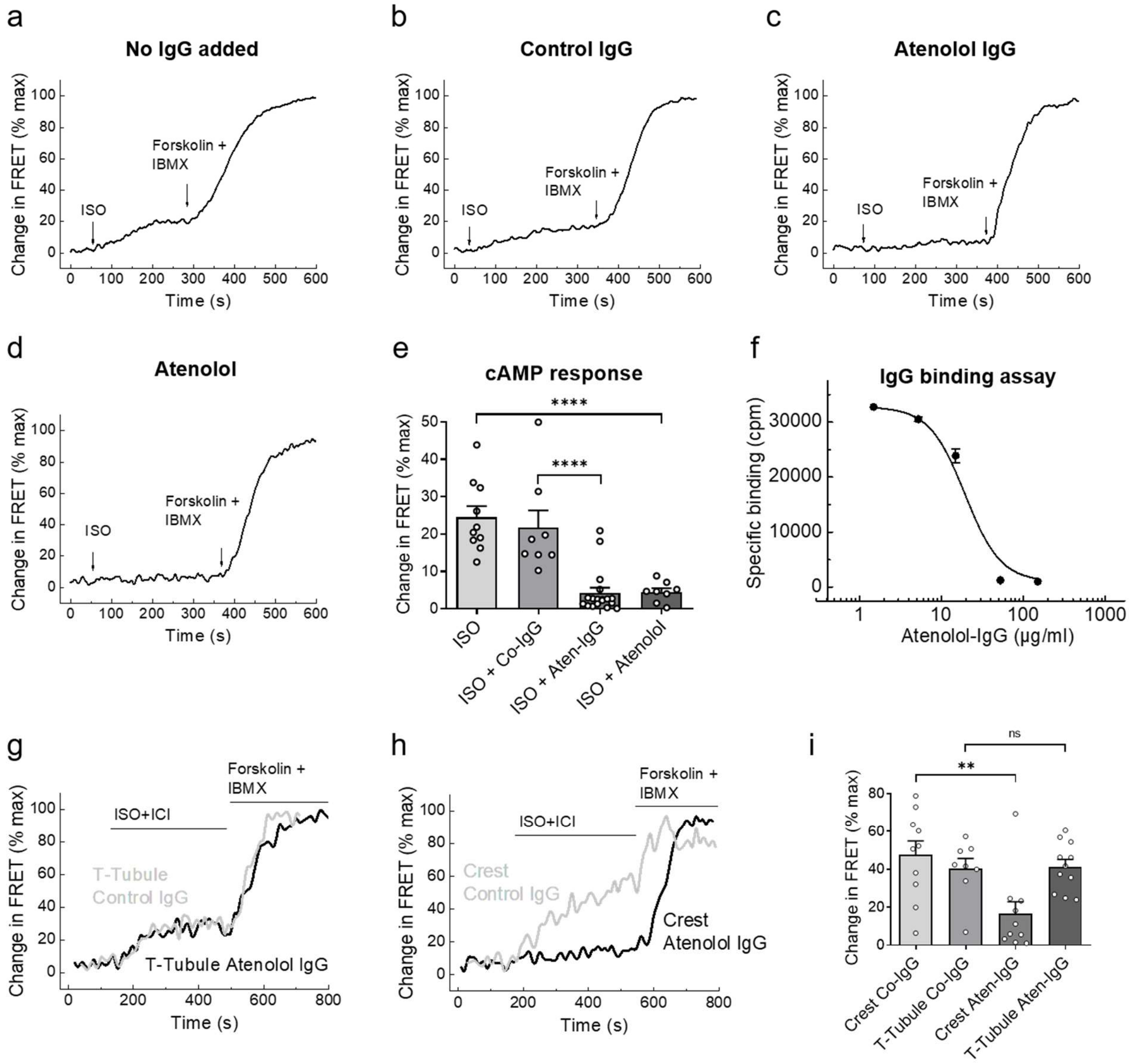


**Extended Figure 7. Beta-blocker activity and receptor binding capacity of Atenolol IgG.** (**a-d**) Representative FRET traces showing cAMP responses to isoproterenol (ISO, 0.03 nM) in HEK293 cells expressing $\beta_1$-AR together with the cytosolic cAMP biosensor Epac1-camps recorded after 30 min pretreatment with vehicle (no IgG added), MSA@IgG (Control IgG, 200 µg/mL, concentration referring to the protein concentration as determined by the Bradford assay) or atenolol@MSA@IgG (Atenolol IgG, 200 µg/mL, concentration referring to the protein concentration as determined with the Bradford assay) or 30 nM of regular atenolol (concentration referring to atenolol; the protein and the atenolol concentrations cannot be directly compared). Maximal sensor response was obtained by stimulation with forskolin (10 µM) and IBMX (100 µM). (**e**) Quantification of FRET responses shown in A-D. n=8-10 cells per condition. **** $p<0.0001$ by one-way ANOVA followed by Sidak multiple comparison test. (**f**) Radioligand binding assay performed on membranes from the HEK293 cells expressing $\beta_1$-AR incubated with the radioligand $^3$H-CGP12177 and increasing concentrations of atenolol@MSA@IgG. Data are mean±SE from 4 independent assays. $IC_{50}$ value for Atenolol-IgG was 21.3±1.7 μg/mL, corresponding to ~140 nM (when calculated using molecular weight of IgG only – $1.5 \cdot 10^5$ g/mol). Compared to the $IC_{50}$ value for atenolol (sFig 4E, 7.7±0.2 μM) this suggests that each Atenolol IgG molecule contains at least ~55 active atenolol molecules when calculated using the molecular weight of the IgG only. In reality, each IgG contains dozens of atenolol-conjugated albumin molecules (each with molecular weight of ~$0.67 \cdot 10^5$ g/mol), so that the real number of atenolol molecules per IgG is 1-2 orders of magnitude greater. (**g-i**) PKA activity in the nucleus was measured using AKAR3-NLS biosensor after local $\beta_1$-AR stimulation by SICM pipette (as described in Figure 1) at T-tubules and cell crests in cell preincubated for 30 min with either Control or Atenolol IgG. Atenolol IgG inhibits only $\beta_1$-AR signals originating from the crest. n=8-11 cells each from 4 rats. n.s. – not significant, ** $p<0.01$ by mixed ANOVA followed by Wald Chi-squared test.

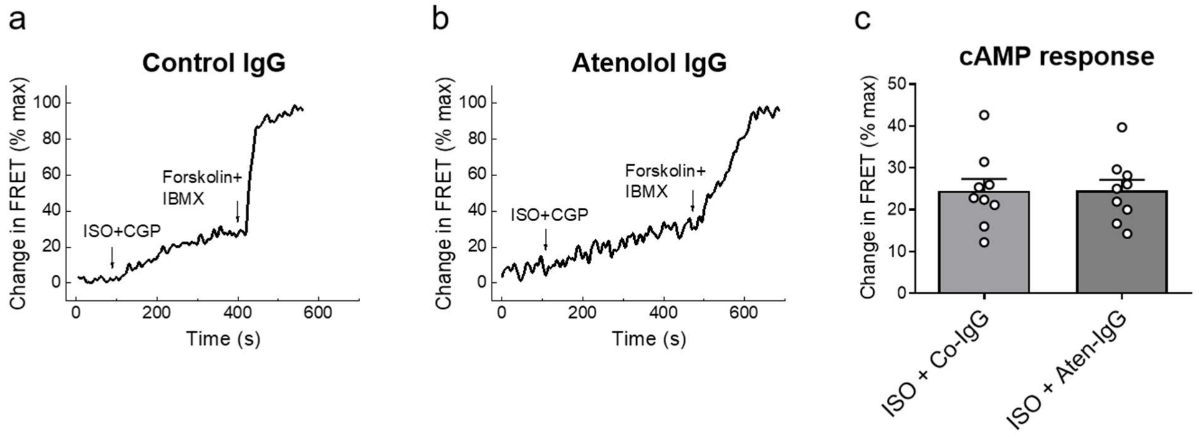


**Extended Figure 8. $\beta_1$-AR selectivity of Atenolol IgG.** (**a,b**) Representative FRET traces showing cAMP responses to isoproterenol (ISO, 0.3 nM) in wildtype HEK293 cells transfected with Epac1-camps endogenously expressing $\beta_2$-AR recorded after 30 min pretreatment with MSA@IgG (Control IgG, 200 µg/mL) or atenolol@MSA@IgG (Atenolol IgG, 200 µg/mL). Maximal sensor response was obtained by stimulation with forskolin (10 µM) and IBMX (100 µM). (**c**) Quantification of FRET responses shown in A,B. n=9 cells per condition. Atenolol IgG does not affect $\beta_2$-AR stimulated cAMP production.

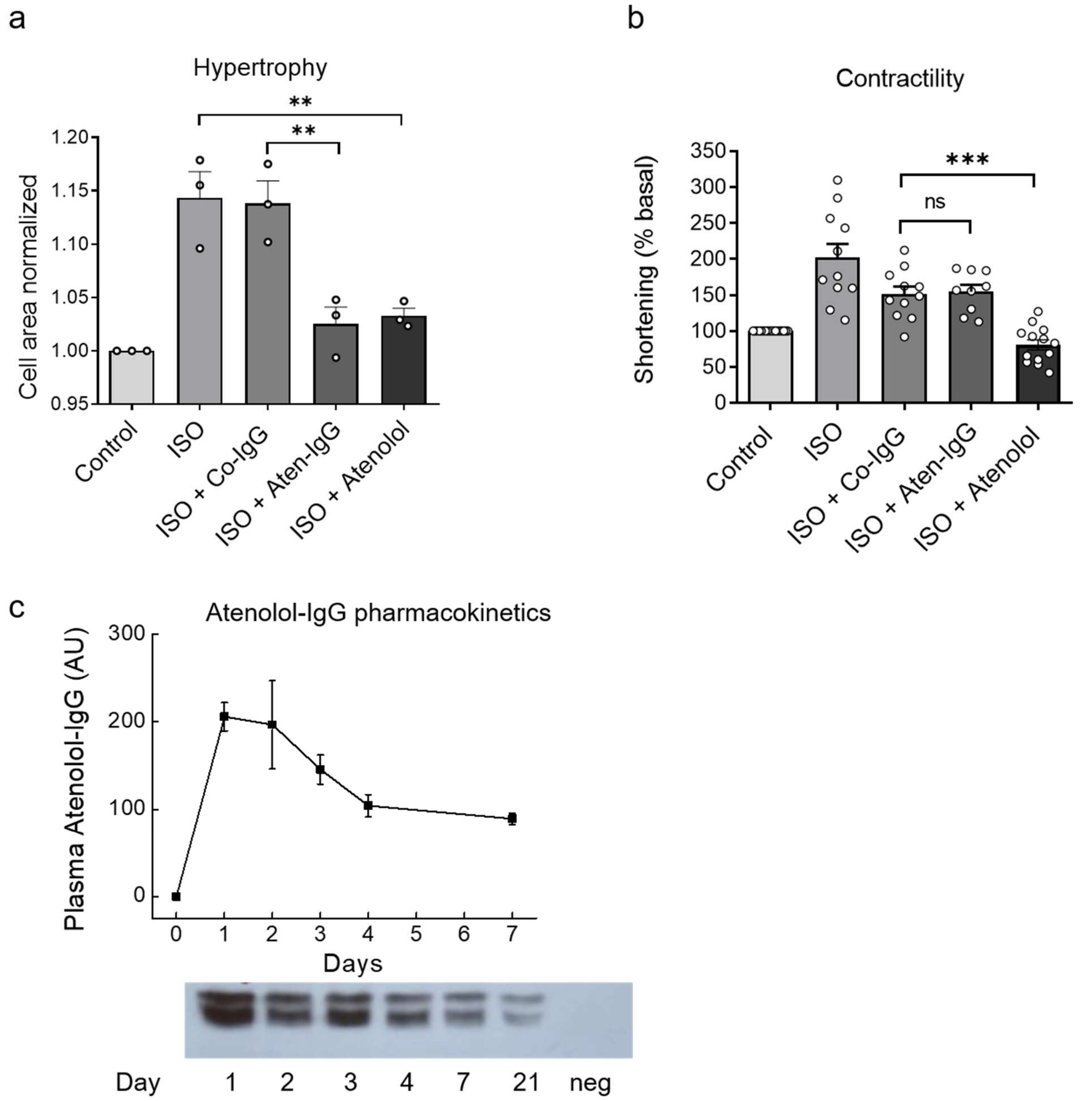


**Extended Figure 9. Atenolol IgG prevents myocyte hypertrophy without contractility decline *in vitro*.** (**a**) Hypertrophy measurement in adult rat myocytes treated for 24 h with 10 nM ISO together with or without atenolol@MSA@IgG (Atenolol IgG), MSA@IgG (Control IgG), both used at 40 µg/mL, or pure atenolol (100 nM). Data are from n=3 independent isolations with at least 100 cells counted per heart. (**b**) Contractility measurements in cells acutely treated with the same compounds (n=9-13 cells each). n.s. – not significant, ** $p<0.01$, *** $p<0.001$ by one-way ANOVA followed by Sidak multiple comparison test. (**c**) Pharmacokinetic analysis of Atenolol-IgG plasma levels at various time points after s.c. injection of 0.6 mg/kg Atenolol-IgG (FITC conjugated for immunoblot detection) shows very long half-life (around one week) of this compound. neg. – negative control with plasma sample from an uninjected animal. Data are means±SE, n=4 each.

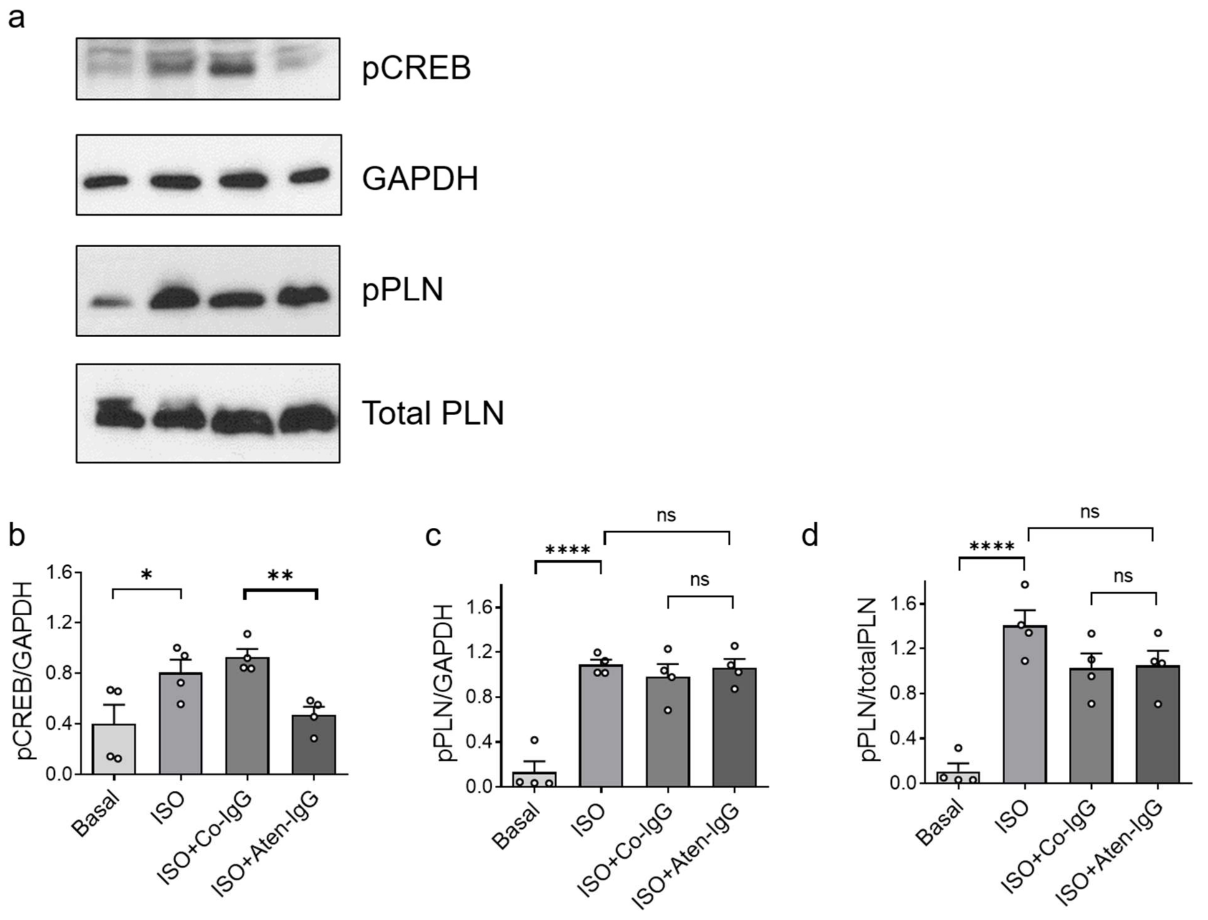


**Extended Figure 10. Atenolol IgG inhibits $\beta_1$-AR induced pCREB without affecting phospholamban (PLN) phosphorylation.** (**a**) Representative immunoblots and quantification of (**b**) pCREB and (**c,d**) pPLN levels in rat cardiomyocytes stimulated for 10 min with 10 nM ISO (in the presence of 50 nM ICI118,551) either without or with 30 min pretreatment with MSA@IgG (Control IgG) or atenolol@MSA@IgG (Atenolol IgG), both at 200 µg/mL. Atenolol IgG inhibit ISO induced CREB but not PLN phosphorylation. GAPDH and total PLN were used as loading control. Data are means±SE, n=4 independent experiments each. n.s. – not significant, * $p<0.05$, ** $p<0.01$, **** $p<0.0001$ by one-way ANOVA followed by Sidak multiple comparison test.

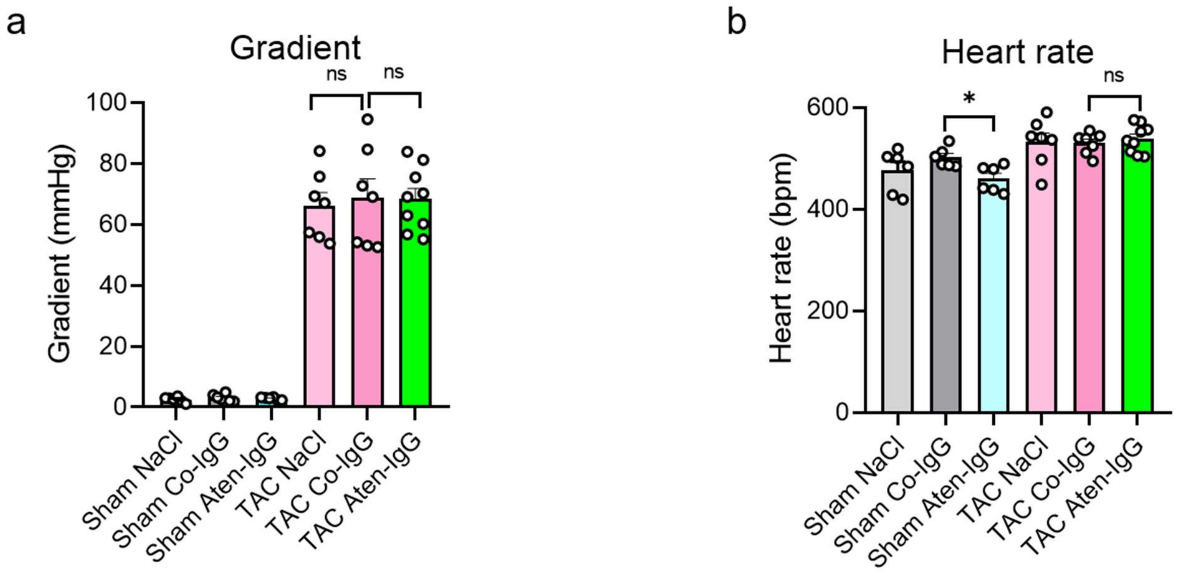


**Extended Figure 11. Echocardiographic parameters measured in sham and TAC mice.** (**a**) Pressure gradients across the constricted aorta and (**b**) heart rates recorded during echocardiography in animals of different treated groups shown in Figure 4E,F. Data are means±SE from n=7-9 mice per group. n.s. – not significant, * $p<0.05$ by one-way ANOVA followed by Sidak multiple comparison test.

**Table S1.**

Characteristics of the heart failure patients used in the study. Values are presented as mean ± SD or n (%). LVEF, left ventricular ejection fraction; LV-EDD, left ventricular end diastolic diameter.

| **Patients, n** | **7** |
|---|---|
| **Male, n (%)** | 4 (66.7) |
| **Female, n (%)** | 3 (33.3) |
| **Myocardial infarction, n (%)** | 3 (33.3) |
| **Diabetes mellitus, n (%)** | 2 (28.6) |
| **Hypertension, n (%)** | 6 (85.7) |
| **Hyperlipidemia, n (%)** | 5 (71.4) |
| **Age at surgery, (years)** | 61 ± 12.9 |
| **Body Mass Index, kg/m²** | 25.4 ± 6.0 |
| **LVEF, %** | 18.3 ± 9.2 |
| **LV-EDD (mm)** | 66.0 ± 2.6 |
